# EZH2/ hSULF1 axis mediates receptor tyrosine kinase signaling to shape cartilage tumor progression

**DOI:** 10.1101/2022.06.07.495151

**Authors:** Zong-Shin Lin, Chiao-Chen Chung, Yu-Chia Liu, Chu-Han Chang, Hui-Chia Liu, Yung-Yi Liang, Teng-Le Huang, Tsung-Ming Chen, Che-Hsin Lee, Chih-Hsin Tang, Ya-Huey Chen, Mien-Chie Hung

**Affiliations:** Graduate Institute of Biomedical Sciences, College of Medicine, China Medical University, Taichung, Taiwan; Center for Molecular Medicine, China Medical University Hospital, Taichung, Taiwan; Department of Biomedical Imaging and Radiological Science, College of Medicine, China Medical University, Taichung, Taiwan; Department of Biotechnology, Asia University, Taichung, Taiwan; Department and Graduate Institute of Aquaculture, National Kaohsiung Marine University, Kaohsiung, Taiwan; Department of Biological Sciences, National Sun Yat-sen University, Kaohsiung, Taiwan

**Author notes:** Correspondence should be addressed to: Mien-Chie Hung, College of Medicine, China Medical University, No. 100, Sec. 1, Jingmao Road, Taichung 406040, Taiwan R.O.C. and Ya-Huey Chen, College of Medicine, China Medical University, No. 91, Hsueh-shih Road, Taichung 40402, Taiwan R.O.C.

## Abstract

**Background:** Chondrosarcomas are primary cancers of cartilaginous tissue and capable of alteration to highly aggressive, metastatic, and treatment-refractory states, leading to a poor prognosis with a five-year survival rate at 11 months for the dedifferentiated subtype. At present, the surgical resection of chondrosarcoma is the only effective treatment, and no other treatment options including targeted therapies, conventional chemotherapies, or immunotherapies are available for these patients.

**Methods:** A non-biased ChIP sequence, cDNA microarray analysis, and validation of chondrosarcoma cell lines identified sulfatase 1(SULF1) as the top EZH2-targeted gene to regulate chondrosarcoma progression. Receptor tyrosine kinase (RTK) array of chondrosarcoma cells with vector control or ectopically expressed SULF1 revealed that cMET was the downstream signal. The regulation of the EZH2/SULF1/cMET axis was further validated in mice and patient samples with mice models and chondrosarcoma tissue array, respectively.

**Results:** The EZH2/SULF1/cMET axis is identified, which contributes to the malignancy of chondrosarcoma and provides a potential therapeutic option for the disease. Ectopically expressed SULF1 or pharmaceutical inhibition of the cMET pathway significantly retards the chondrosarcoma growth and extends mice survival.

**Conclusions:** The results not only established a signal pathway promoting the malignancy of chondrosarcoma but also provided a therapeutic potential for further development of effective target therapy to treat chondrosarcoma.

## Introduction

Chondrosarcoma, a malignant cartilaginous tumor, is the most common primary skeletal malignancy affecting bone (Fletcher, 2002). Accumulation of mutations in molecules regulating cell growth control, programmed cell death and DNA instability is required for transformation from cartilage neoplasia to malignancy tumors. They are further classified into primary central and secondary peripheral chondrosarcomas based on their location in the bone (Gelderblom et al., 2008). The most common sites are bones of the pelvis, followed by the diaphysis or metaphysis of the proximal femur and humerus, distal fumer, and ribs. The histologies in central and peripheral chondrosarcomas are similar, and all three different grades are discerned. They are the best predictor of clinical behavior at present (Evans, Ayala, & Romsdahl, 1977). Metastasis is rare in Grade I chondrosarcomas while metastases occur in 10% of Grade II and 71% of Grade III chondrosarcomas (Bjornsson, McLeod, Unni, Ilstrup, & Pritchard, 1998). Conventional chondrosarcomas are extremely resistant to chemotherapy and radiotherapy, resulting in limited treatment options (Choy et al., 2012; Gelderblom et al., 2008). Depending on the grade of the chondrosarcoma, a therapeutic surgical approach remains the only practical treatment (Lee et al., 1999). After surgery, patients with poor-quality life and for those inoperable patients (70 %), radiation therapy alone is an option with uncertainty outcome (Fromm et al., 2018). Additionally, the relatively poor vascularity results in more difficult delivering drugs than to other vascularized cancers (Bovee, Hogendoorn, Wunder, & Alman, 2010; Gelderblom et al., 2008; Lee et al., 1999). Due to rarity and various subtypes of chondrosarcoma, lack of interest from pharmaceutical industry obstructs the drug development and conduct of clinical trials for these patients (Miwa et al., 2019). Until now, there are no effective treatments for patients with inoperable or metastatic disease (70 %), and therefore, it is an un-met medical need to develop new treatment modalities. IGF inhibitors have been used to treat chondrosarcomas by increasing the rate of apoptosis (Ho et al., 2009; Olmos et al., 2010). Small molecule tyrosine kinase inhibitor dasatinib is known to reduce cell viability in chondrosarcoma cell lines by targeting SRC (Schrage et al., 2009). Recently, several target therapies were applied in phase I or II clinical trials in chondrosarcoma, including isocitrate dehydrogenase (IDH) inhibitor, tyrosine kinase inhibitors targeting vascular endothelial growth factor receptor (VEGFR), platelet derived growth factor receptor (PDGFR), insulin-like growth factor receptor (IGF-1R)(Truong, Cherradi-Lamhamedi, & Ludwig, 2022). These clinical trials are encouraging, however remain in the early stage and the clinical outcomes are yet to observe.

Enhancer of Zeste 2 (EZH2) is the major component of the polycomb repressor complex 2 (PRC2) (Kuzmichev, Nishioka, Erdjument-Bromage, Tempst, & Reinberg, 2002). PRC2 has been reported to repress gene expression through the histone methyltransferase EZH2-catalyzed trimethylation of histone 3 lysine 27 (H3K27) to establish repressive epigenetic marker (Cao et al., 2002; Cao & Zhang, 2004; Kuzmichev et al., 2002). EZH2 is required to maintain DNA methylation and suppress specific genes such as tumor suppressor genes by direct interaction of EZH2 and D NA methyltransferases (DNMT1, DNMT3A, and DNMT3B) (Sauvageau & Sauvageau, 2010). EZH2 possesses the catalytic ability to transfer a methyl group from S-adenosyl methionine (SAM) to H3K27 in the PRC2 and represses the expression of target genes including a broad range of genes such as those in the regulation of cell cycle, DNA damage repair, cell fate and differentiation, senescence, apoptosis, and cancer (Cao et al., 2002). Overexpression of EZH2 has been observed in lymphomas, sarcomas, and cancers of the breast, colon, liver, lung, and prostate (Chang & Hung, 2012; Han et al., 2020; X. Li et al., 2009; Sauvageau & Sauvageau, 2010; Yamaguchi & Hung, 2014). It has been reported that EZH2 mediates different types of human cancer through transcriptional, post-transcriptional, and post-translational stages, demonstrating that transcriptional factors, such as E2F, are able to bind to the promoters of EZH2 and EED, which is necessary for E2F modulated-cell proliferation (Bracken et al., 2003; Wei et al., 2011).

EWS-FLI1, a fusion oncoprotein in Ewing’s sarcoma, can transcriptionally activate EZH2 expression, and upregulated EZH2 expression plays a vital role in tumor growth and endothelial/ neuroectodermal differentiation (Richter et al., 2009). More recently, Taniguchi et al. demonstrated that EZH2 directly targets the Kruppel-like factor 2 (KLF2), which is a tumor suppressor, and interrupts its functions in cell cycle arrest and apoptosis, leading to tumorigenesis in the osteosarcoma mouse model (Taniguchi et al., 2012). In addition, a number of post-translational modifications also regulate EZH2-mediated signaling (Cha et al., 2005; Chang et al., 2011; Nie et al., 2019; Wan et al., 2018; Wei et al., 2011). These studies all point to a crucial role of EZH2 in tumorigenesis through a number of events in different human cancers including sarcoma. Notably, the dysregulation of differentiation programs of mesenchymal stem cells (MSCs) may lead to osteosarcomas and chondrosarcomas (Mohseny & Hogendoorn, 2011). Our previous results demonstrated that EZH2 might be a key modulator in bone development and sarcoma pathogenesis (Chen et al., 2016; Wei et al., 2011).

Human sulfatase 1 (SULF1) was characterized and revealed to desulfate cellular heparin sulfate proteoglycans (HSPGs). Sulfated HSPGs play a vital regulator of heparin binding growth factor signaling: e.g., FGF2, VEGF, HGF, PDGF and heparin binding EGF (HB-EGF), also known as receptor tyrosine kinase (RTK) signaling (J. Lai et al., 2003). Therefore, enzymes that desulfate HSPGs could have a tumor suppressor effect via abolishment of ligand-receptor binding and downstream signaling. Complete loss or markedly attenuated expression of SULF1 appeared in cancer cell lines derived from breast, pancreas, kidney and hepatocellular cancers, suggesting that down regulation of SULF1 is relatively common among epithelial cancers (J. P. Lai, Sandhu, Shire, & Roberts, 2008).

In this study, we integrated EZH2-ChIP sequence and cDNA microarray analysis then validated in multiple chondrosarcoma cell lines to identify SULF1 as a top EZH2-targeted gene governing chondrosarcoma progression. We unravel the critical roles of SULF1 in suppressing tumor growth of chondrosarcoma by reducing RTK signaling pathway such as cMET which is known to increase malignancy and causes resistance to target therapy (Chu et al., 2020; Du et al., 2016; H. Li et al., 2019; Sun et al., 2020). Using pharmaceutical inhibitors targeting at EZH2 or cMET (Yang et al., 2015) to treat chondrosarcoma mice model significantly reduced tumor growth and prolonged mice survival. The results offer an opportunity for the development of effective target therapy to treat chondrosarcoma.

## Results

### EZH2 may have a role in chondrosarcoma development

Overexpression of EZH2 has been observed in sarcomas, lymphomas and cancers of the breast, colon, liver, lung, and prostate (Chang & Hung, 2012; Sauvageau & Sauvageau, 2010). The abnormal differentiation programs of MSCs regulated by EZH2 results in osteosarcomas and chondrosarcomas (Mohseny & Hogendoorn, 2011). In addition, EZH2 was shown to govern bone development (Chen et al., 2016) and sarcoma pathogenesis (Wei et al., 2011). Thus, we hypothesize that EZH2 might have a role in the chondrosarcoma.

To examine the potential role of EZH2 in chondrosarcoma malignancy, we initially conducted western blot to analyze the expression of EZH2 in normal chondrocytes and chondrosarcoma cell lines. In contrast to normal chondrocytes, EZH2 was highly expressed in CH2879 and JJ012 chondrosarcoma cell lines as well as increased histone 3 trimethylation at lysine 27 (H3K27me3) (Fig. 1A). Due to lack of specific chondrosarcoma database of TCGA, we further systematically examined the prognostic significance of EZH2 in sarcoma, a group including chondrosarcoma, using the median values for patient stratification analyzed by the Kaplan-Meier Plotter, a tool for meta-analysis-based biomarker assessment (Fig. 1B). Consistently, the expression of EZH2 correlated well with poor survival of sarcoma patients and the justification of clinical relevance for EZH2 in chondrosarcoma by using tissue microarray was also shown later in Fig. 8.

**Fig 1.**
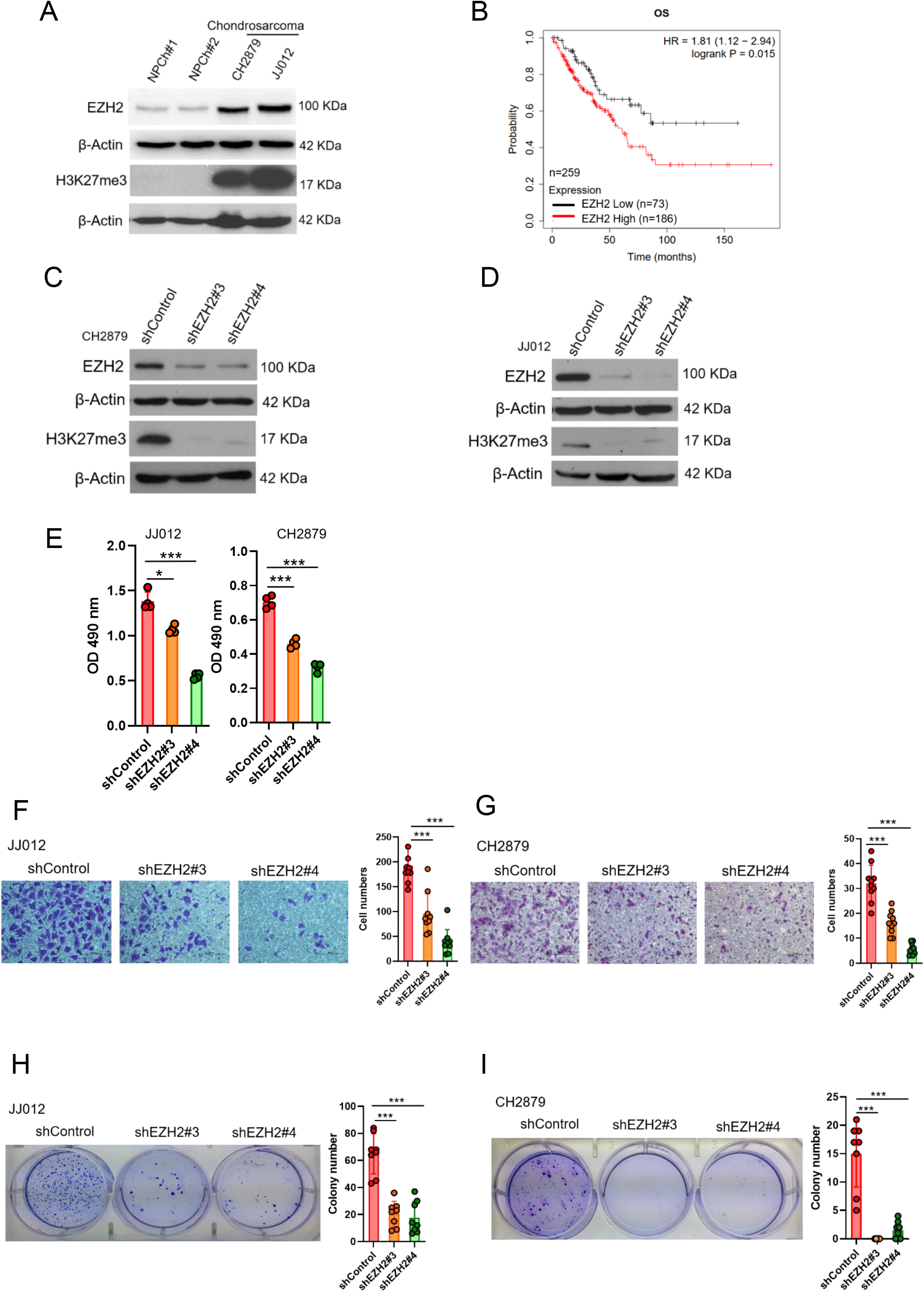

To validate the role of EZH2 in tumorigenicity of chondrosarcoma, we introduced shEZH2 or control shRNA by lentiviral system into chondrosarcoma cell lines (Fig. 1C, 1D). Knockdown of EZH2 reduced the proliferative, migrated and colony-forming capacity of chondrosarcoma cell lines (Fig. 1E, 1I) and, supporting the notion that EZH2 may have a role in chondrosarcoma development.

### SULF1 is an EZH2-targeted gene and downregulated in chondrosarcoma

To identify EZH2-targeted genes in chondrosarcoma, we used cDNA microarray and EZH2-chromatin immunoprecipitation sequencing (ChIP-seq) profiling to identify the dysregulated genes which were targeted by EZH2 in chondrosarcoma cell lines. We first searched the genes that were both targeted by EZH2 (n=704) and associated with significant dysregulated genes from cDNA microarray in comparison with normal chondrocyte to JJ012 chondrosarcoma cell line (n=6771). Between these two pooled genes, 40 were overlapped. Because EZH2 is a gene silencer and catalyzes H3K27 trimethylation (H3K27me3) in the nucleus to modulate chromatin compaction followed by transcriptional suppression of downstream genes. Thus, we hypothesized that upregulated-EZH2 may silence tumor-suppressive genes. To further reduce the number of candidate regulators, we first focused on genes function as a tumor suppressor role among those downregulated by EZH2. This analysis resulted in 7 known candidate genes (Fig. 2A).

**Fig 2.**
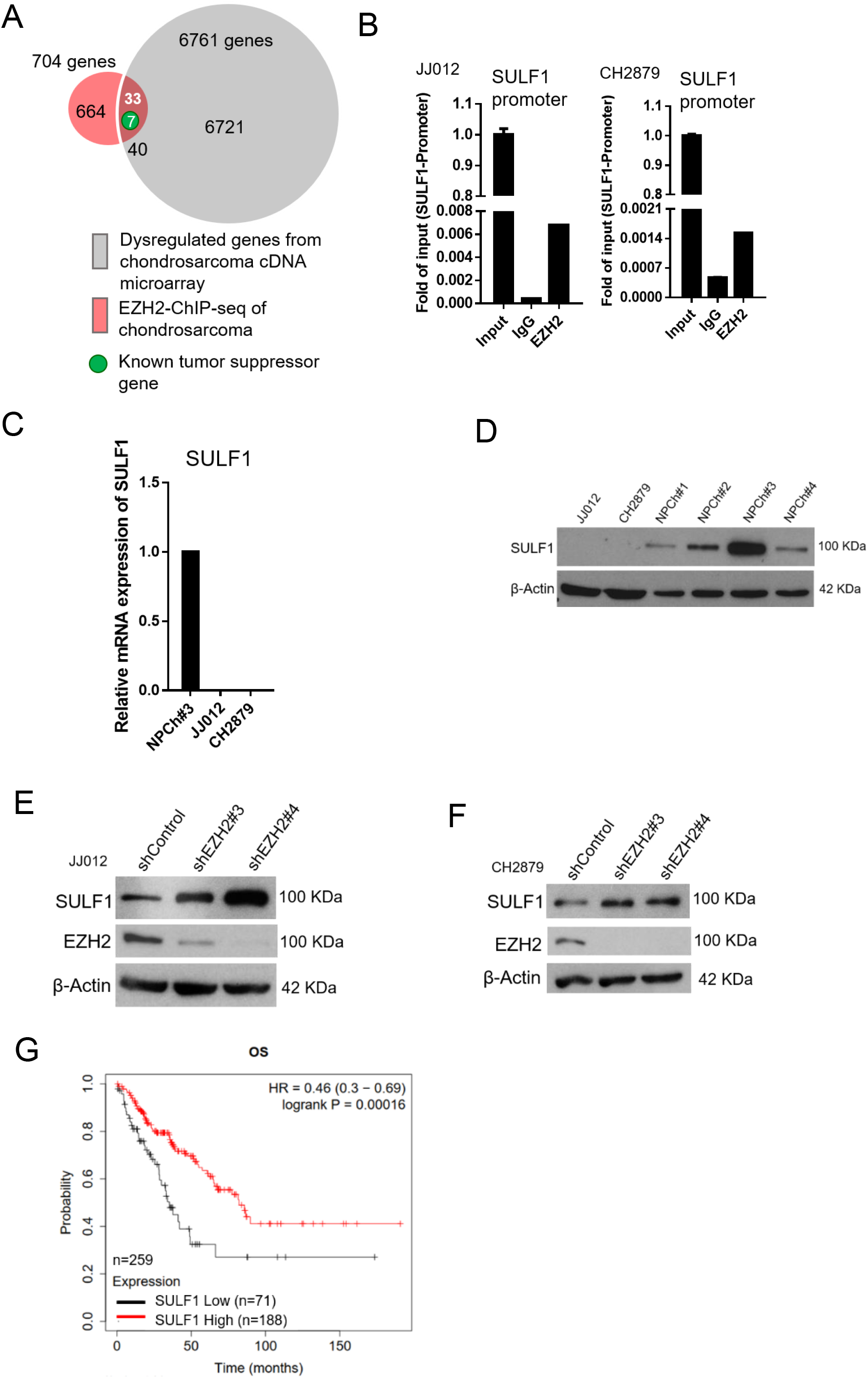

Of note, human sulftase1 (SULF1) was the top among those reduced expression in chondrosarcoma by EZH2 targeting. q-ChIP experiments further validated enrichment of EZH2 bound at Sulf1 promoter in various chondrosarcoma cell lines (Fig. 2B). Consistently, qRT-PCR analysis showed that Sulf1 was down-regulated in chondrosarcoma cell lines, JJ012 and CH2879 as compared with normal chondrocyte (Fig. 2C). Immunoblotting also demonstrated that SULF1 expression was reduced in JJ012 and CH2879 cell lines (Fig. 2D). Knockdown of EZH2 induced SULF1 expression in both JJ012 and CH2879 cell lines (Fig. 2E, 2F). The results indicated that SULF1 is the direct targeted gene of EZH2 in chondrosarcoma cell lines. This regulation of EZH2/ SULF1axis was also observed in the other sarcoma cells such as osteosarcoma cell lines (Supplementary Fig 1A-1G). To further investigate the prognostic significance of EZH2/ SULF1 axis in sarcoma, the Kaplan-Meier Plotter indicated that lower SULF1 expression significantly correlated with poorer survival in sarcoma populations (Fig. 2G).

### Ectopic expressed SULF1 attenuates progression of chondrosarcoma

To further explore the potential effect of SULF1 on the chondrosarcoma progression, stably expressed SULF1 of chondrosarcoma cell lines were established (Fig. 3A) and assays for cell proliferation, migration, and colony-forming ability were conducted. SULF1 ectopic-expression inhibited chondrosarcoma cell proliferation (Fig. 3B), migration (Fig. 3C), and colony-forming capacity (Fig. 3D). Chondrosarcoma cells with ectopic-expressed SULF1 were re-introduced control and specific SULF1 shRNA and followed by the test of colony-forming assay. The colony-forming capacity was restored in the presence of SULF1 shRNA as compared with shRNA control (Fig. 3E). We also examined the role of SULF1 in chondrosarcoma growth using an alternative tumor xenograft model. Subcutaneous injection of CH2879 cells resulted in reliable tumor growth in nude mice. The stable transfection of CH2879 cells with SULF1 markedly repressed the growth of chondrosarcoma tumors with smaller size and weight compared with the control counterparts (Fig. 3F). It is worth to mention that the SULF1 level in the transfected chondrosarcoma cell lines are comparable to the expression level of SULF1 in primary normal chondrocytes (n=107) by using qRT-PCR analysis (Supplementary Fig. 2). All these results revealed that SULF1 is a downstream target of EZH2 and serves as a tumor suppressor role in chondrosarcoma development.

**Fig 3.**
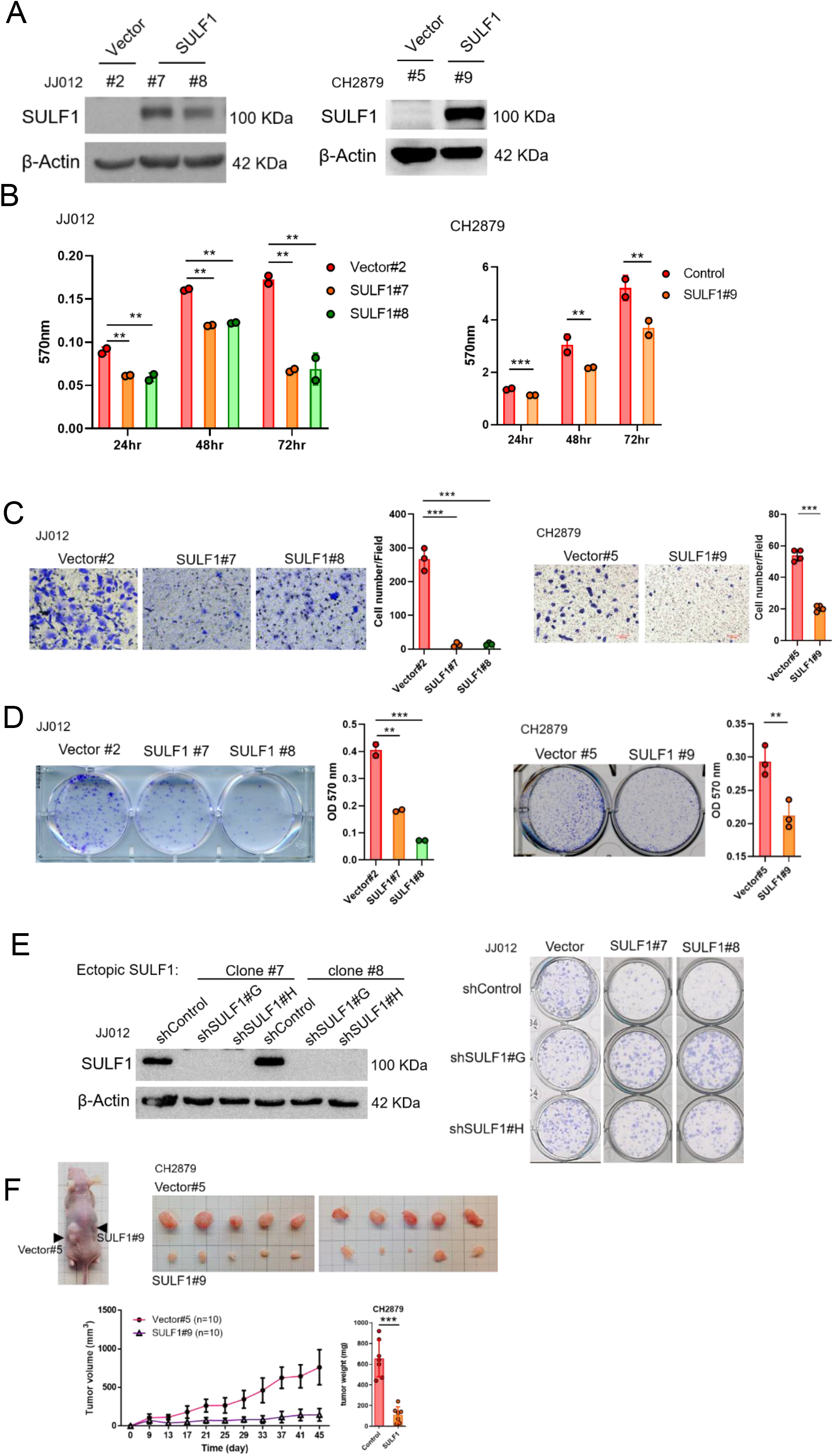

### SULF1 repressed the signaling pathway of cMET via dissociation between of cMET and its co-receptor (HSPG) in chondrosarcoma

To understand the mechanism of SULF1-mediated inhibitory effects on the tumorigenesis of chondrosarcoma cell lines, the unbiased RTK antibody array was performed and revealed that several phosphorylation of RTKs were decreased in SULF1 overexpression cell lines, with the largest fold decreases in cMET, and followed by ALK and TIE2 (Fig. 4A, 4B). To further confirm the phosphorylation of cMET in vector and SULF1 stably expressed chondrosarcoma cell line from antibody array, western blotting analysis was conducted. Furthermore, downstream signaling of cMET like AKT, ERK1/ 2 and p38 were also tested. Western blotting analysis indicated that in cells with ectopic expression of SULF1, phosphorylation of cMET, AKT, ERK1/ 2 and p38 was reduced (Fig. 4C).

**Fig 4.**
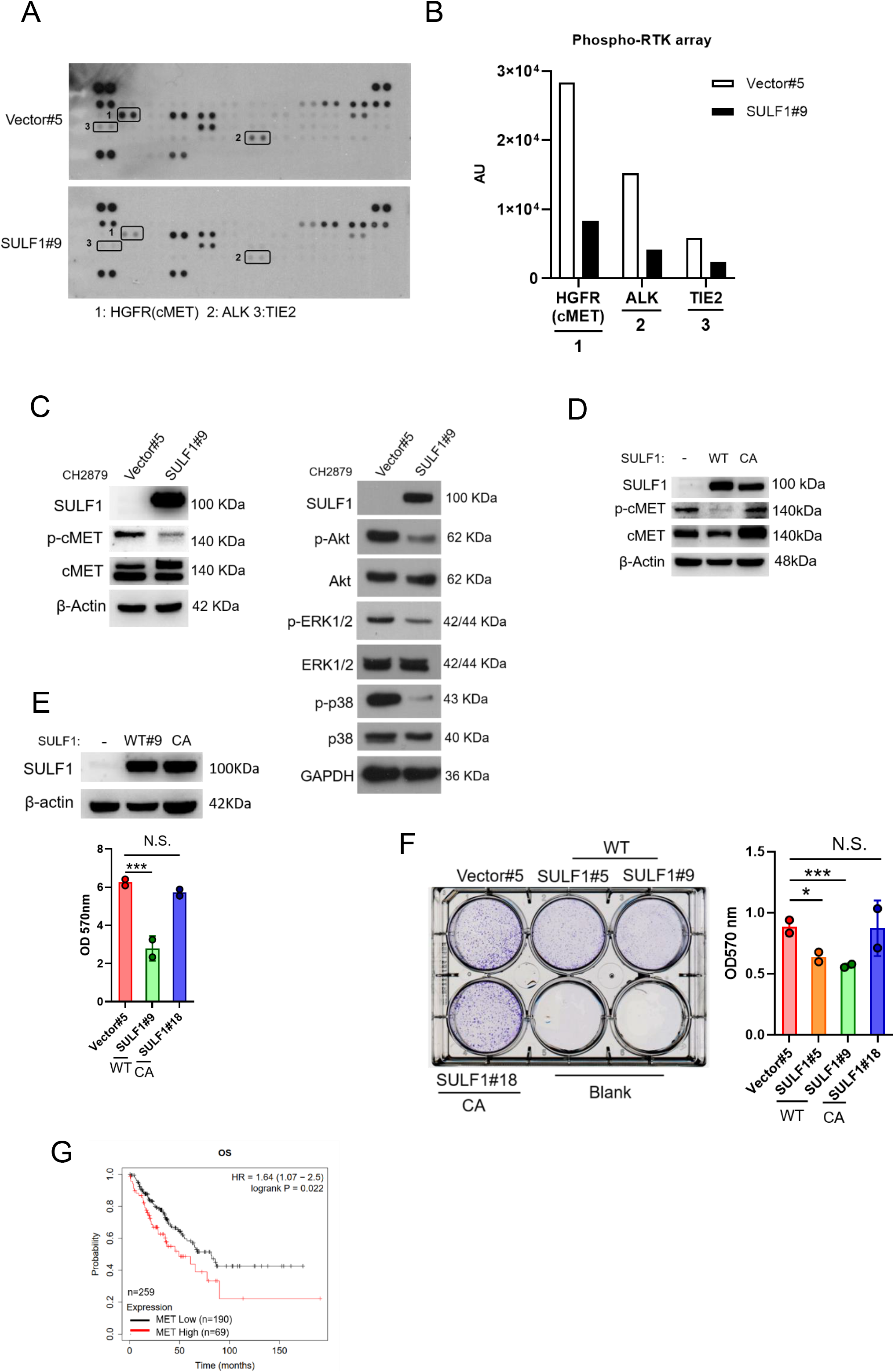

Prior investigation reported that the SULF1 contain a large hydrophilic domain (HD), located between the N-terminal catalytic domain and C-terminal domain. The function of the HD of human SULF1 in governing enzyme activity, cell surface targeting, secretion, and substrate recognition. The double mutant SULF1 C87A/ C88A was completely inactive (Frese, Milz, Dick, Lamanna, & Dierks, 2009), serve as an enzymatically inactive SULF1 C87A/ C88A (CA). To explore whether the phosphorylation of cMET is mediated by SULF1 enzyme activity, we generated stably expressed either wild type or CA mutant of SULF1 in CH2879 cell line. The CA mutant of SULF1 lost enzymatic activity and was unable to repress the phosphorylation of cMET in compared to the ectopic expressed wild type SULF1 in CH2879 (Fig. 4D), suggesting that sulfatase enzymatic activity of SULF1 was required for the dephosphorylation of cMET.

Overexpression of wild type SULF1 reduced the proliferation (Fig. 4E) and colony forming ability (Fig. 4F), as expected from the reduced phosphorylation of cMET in Fig. 4D. In contrast, CA mutant SULF1 which lost enzymatic activity failed to do so in chondrosarcoma cell line. Additionally, higher cMET expression associated with poorer patient survival was also observed (Fig. 4G). Taken together, overexpression of wild type SULF1 is implicated conferring decreased phosphorylation of cMET to attenuate cancer related behavior in chondrosarcoma cell lines, and the enzyme activity of SULF1 is required for this activity.

Heparan sulfate proteoglycans (HSPGs) act as co-receptors in cell signaling pathways and provide binding sites for growth factors and morphogens via specific sulfation patterns on their heparan sulfate (HS) chains. Through enzymatically removing 6-O-sulfate groups from HSPGs on the cell surface by SULF1, it can regulate the tricomplex formation among RTK, HSPGs and ligands, thereby modulating important processes such as development, cell growth, and differentiation (J. P. Lai et al., 2008). A high affinity ligand of cMET, hepatocyte growth factor/ scatter factor (HGF/ SF) is well known to stimulate cMET activity and trigger important cellular processes including to enhance cell proliferation, invasion, survival and angiogenesis. The HSPGs can bind to various growth factors and function as signal co-receptors. Actually, CD44 allows to increase the local concentration of glycosaminoglycan-associating proteins, such as osteopontin (OPN), FGF2, VEGF and HGF, thereby inducing the capacity of these ligands to interact with their receptors and so reducing the threshold for signal transduction (A. Z. Lai, Abella, & Park, 2009; Thayaparan, Spicer, & Maher, 2016). Interaction of CD44 and cMET facilitated tumor cell migration, invasion and metastasis (Jeon & Lee, 2017; Zoller, 2011). Most of these interactions induce synergistic effects on cancer progression and the generation of resistance to therapy (Engelman et al., 2007; Viticchie & Muller, 2015).

Thus, we hypothesized that SULF1 may modulate cMET signaling by desulfating cell surface heparan sulfate glycosaminoglycans (HSGAGs) and thus abrogating HSGAG-dependent growth signaling. A model was shown in figure 8D.

To further test this hypothesis, the phosphorylation of cMET was examined by western blotting in vector and ectopic SULF1 expressed chondrosarcoma cell lines treated with or without HGF ligand. The phosphorylation of cMET was dampened in SULF1 overexpressed chondrosarcoma cell lines (Fig. 5A, 5B). The interaction between cMET and CD44 was clearly detected by immunoprecipitation analysis in vector control chondrosarcoma cell lines as reported, yet such interaction between cMET and CD44 was abolished in SULF1 overexpressed chondrosarcoma cell lines (Fig. 5C). To clarify whether SULF1 expression leads to desulfation of cell surface HSGAG, we performed flowcytometry and western blot utilizing the 10E4 anti-HSPG antibody to recognizes N-sulfated glucosamine residues. The ectopic expressed SULF1 of chondrosarcoma cell lines showed the cell surface staining for N-sulfated glucosamine containing HSGAGs was diminished in compared with vector control chondrosarcoma cell lines via using flowcytometry analysis (Fig. 5D, 5E). Furthermore, western blot of sulfated HSGAGs in SULF1 ectopic expressed cell line was reduced in contrast to the counterpart (Fig. 5F). Together, these results indicated that downregulation of SULF1 enhances the complex formation of cMET, HGF, and CD44, consequently to increase the downstream survival signaling to promote tumorigenesis of chondrosarcoma.

**Fig 5.**
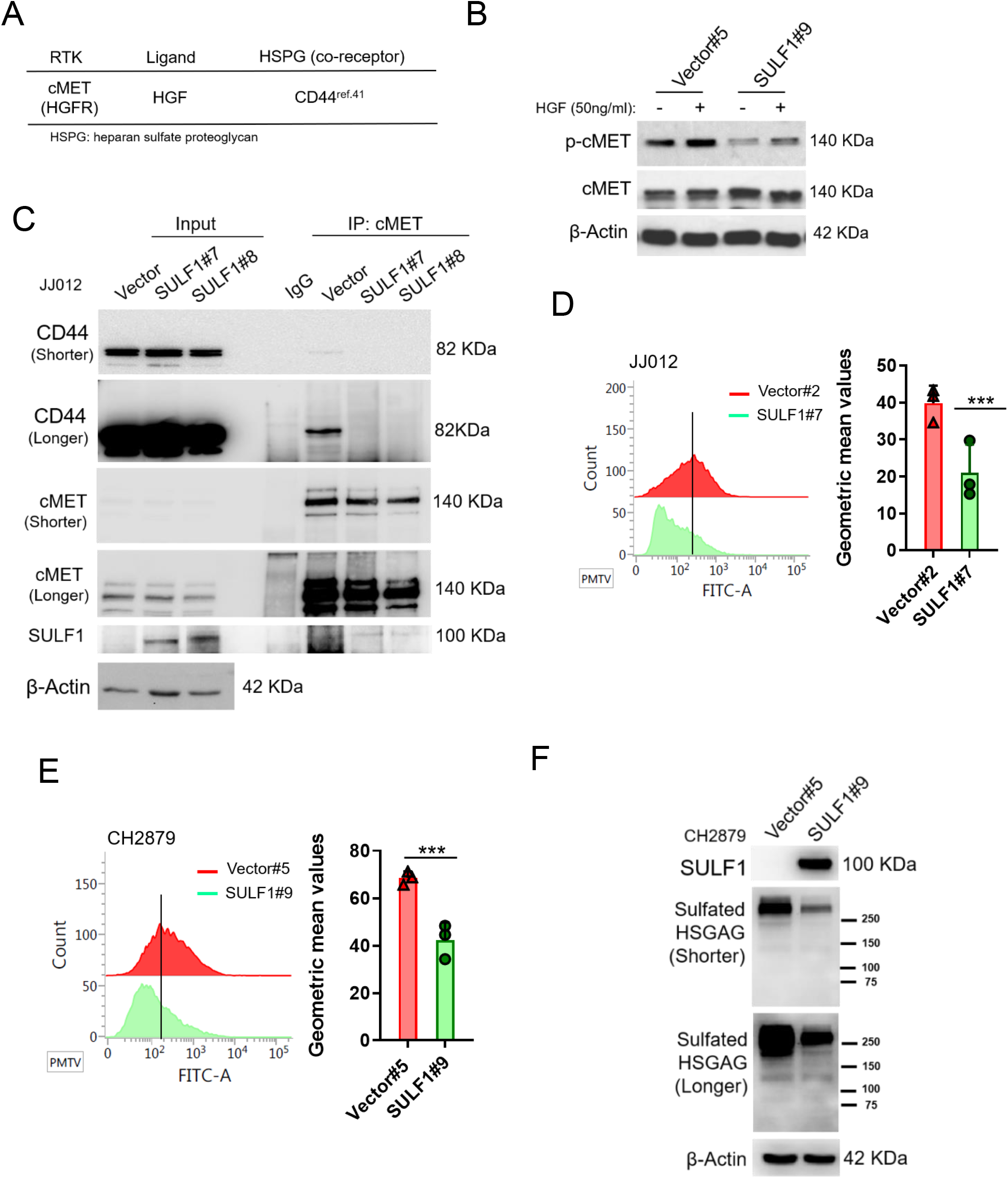

### Treatment with cMET pharmaceutical inhibitors effectively blocks chondrosarcoma growth and prolongs mice survival

We, therefore, determined whether pharmacological inhibition of EZH2 or cMET activity could be an option for target therapy for chondrosarcoma. EZH2 inhibitor such as GSK343 and tazemetostat, which compete with Sadenosyl-methionine for binding to EZH2, thereby inhibiting histone methyltransferase activity without affecting EZH2 protein expression was examined. The cMET small molecule inhibitors, tivantinib, capmatinib and crizotinib, were also used to test the treatment efficacy. In contrast to cMET inhibitors, EZH2 inhibitor required higher IC50 concentrations (Supplementary Fig. 3A, 3B) to diminish trimethylation of histone lysine 27 (H3K27me3) (Supplementary Fig. 3C) and colony-forming ability (Supplementary Fig. 3D, 3E) in chondrosarcoma cell lines. For cMET inhibitors, the IC50 of campatinib (12 μM) was much higher than tivantinib (1 or 1.2 μM) and crizotinib (0.5 μM) in chondrosarcoma cell lines (Supplementary Fig. 4A, 4B). Tivantinib, crizotinib and capmatinib effectively attenuated the phosphorylation of cMET in both CH2879 and JJ012 chondrosarcoma cell lines (Fig. 6A). Particularly, tivantinib and crizotinib decreased the phosphorylation of cMET with low dosage at 0.4 and 0.3 μM, respectively (Fig. 6B). Thus, we focused on tivantinib and crizotinib inhibitors for the further assays. Reduction of colony forming capacity was also observed in both CH2879 and JJ012 cell lines by treated with tivantinib and crizotinib (Fig 6C, 6D). Collectively, the treatment efficacy of cMET inhibitors are effectively to reduce colony formation in chondrosarcoma cell lines.

**Fig 6.**
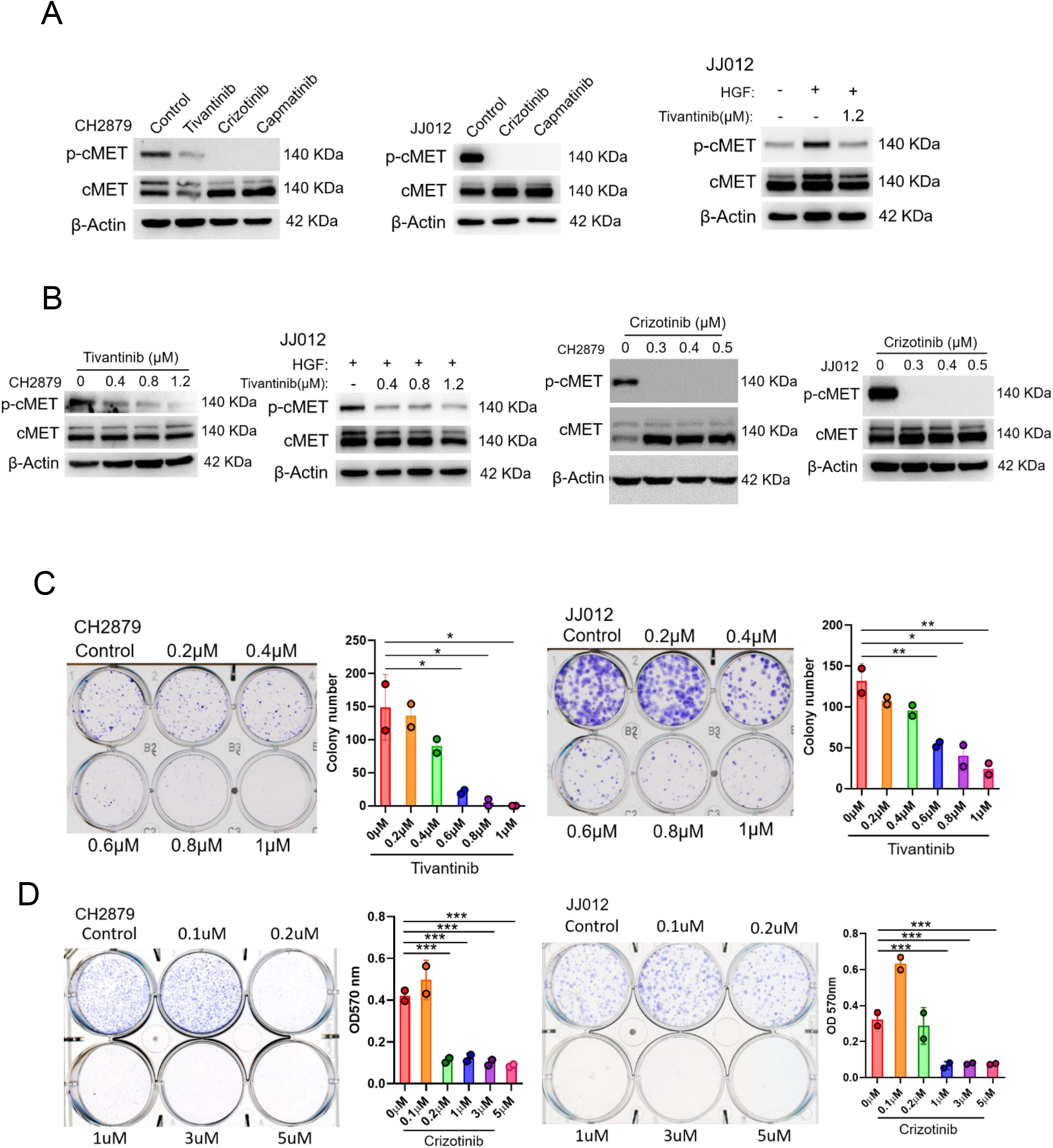

Next, we explored the effects of pharmacologically targeting cMET on tumor growth in mice. Mice with established luciferase-CH2879 tumors were treated with vehicle or crizotinib (Fig 7A). Crizotinib treatment inhibited cartilage tumor growth in orthotopic immunodeficiency mice in a dose-dependent manner (Fig. 7B, 7C) without altered the mice body weight during administrated vehicle or crizotinib (Fig. 7D). Moreover, western blotting was shown that crizotinib therapy could decrease the phosphorylation of cMET in mice tumors (Fig. 7F), which results in attenuating chondrosarcoma progression and dramatically ameliorate survival (Fig. 7G). Overall, these results demonstrated that crizotinib had therapeutic implications to inhibit the growth of chondrosarcoma in mice.

**Fig 7.**
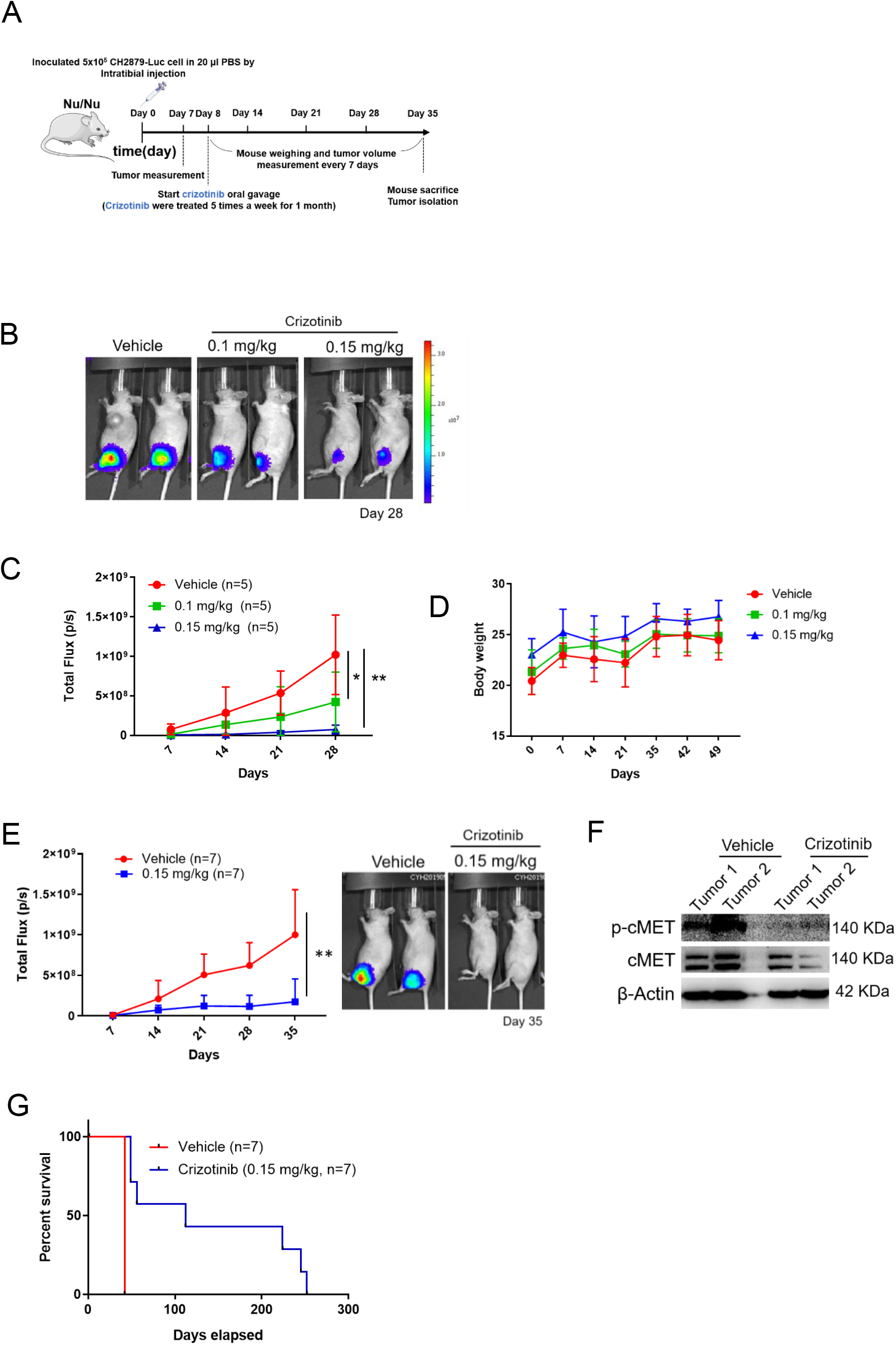

### Clinical significance of the SULF1/ cMET pathway in chondrosarcoma

To further validate our observation in clinical setting, we examined the clinical significance of EZH2, SULF1, and phospho-cMET by analyzing H-score of immunohistochemistry (IHC) on commercially available human bone cancer tissue array. As a result, the patient samples exhibited a low-level expression of SULF1 and high-level expression of EZH2, and phospho-cMET in compared to their counterpart (Fig. 8A-8C). These similar results were also observed in osteosarcoma tissue samples (n=31) (Supplementary Fig. 5). Together, our data demonstrate that EZH2-mediated downregulation of SULF1 are critical for de-suppressed cMET signaling pathway in bone cancer. In chondrosarcoma, the expression of EZH2 was upregulated and thereby inhibited the SULF1 expression. Reduction of SULF1 cannot remove the 6-O sulfation in HSPG resulting in the stabilization of HGF, cMET and CD44 tricomplex to activate the downstream signaling such as phosphorylation of AKT, p38, and ERK (Fig. 8D).

**Fig 8.**
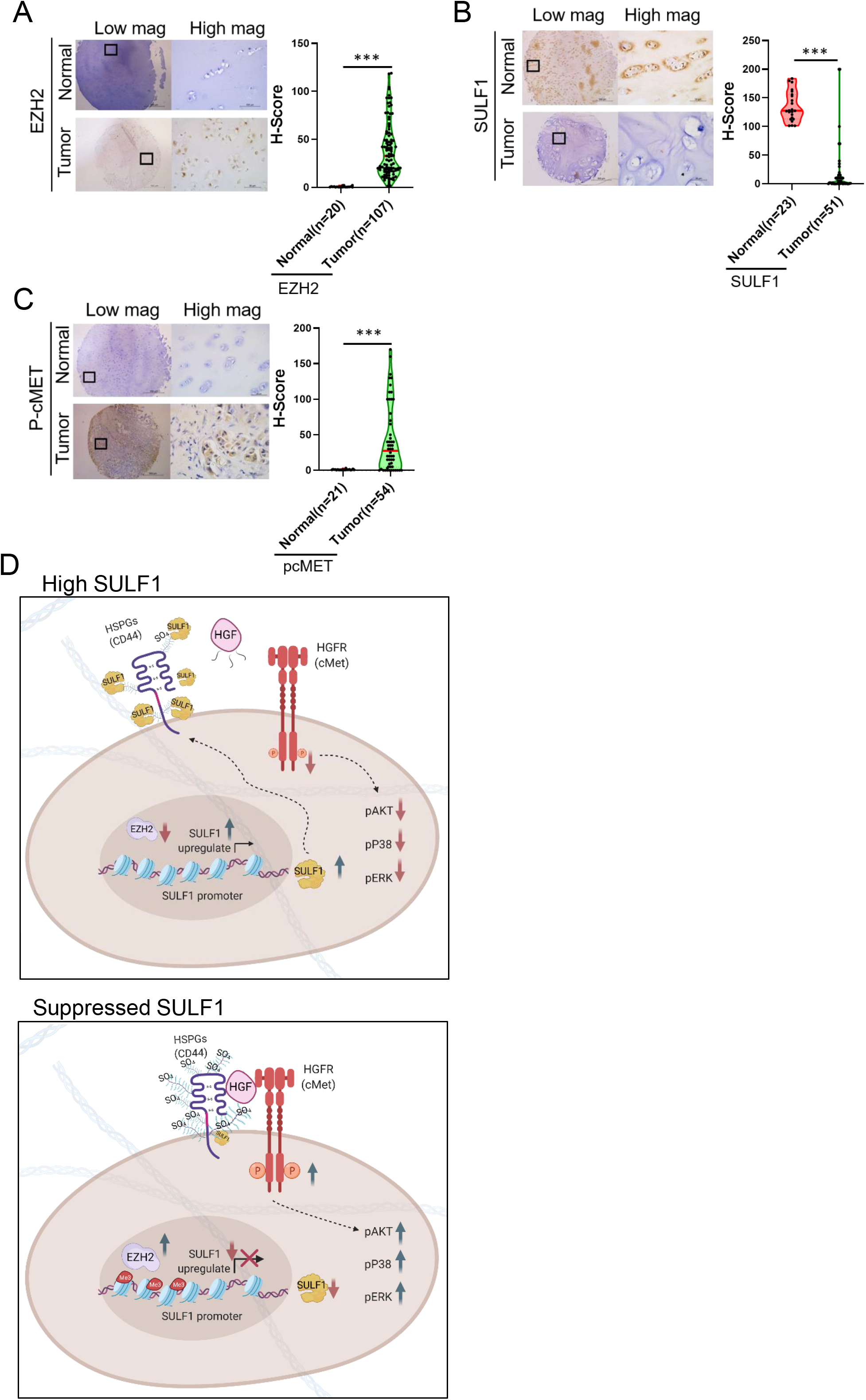

## Discussion

Chondrosarcomas are heterogeneous group of malignant cartilaginous neoplasms with various morphological features, represented by resistance to chemo-and radiation therapies. Mounting molecular alteration involved in chondrosarcoma pathogenesis have characterized in the last several decades (Amary et al., 2011; Tarpey et al., 2013; Totoki et al., 2014; Zhang et al., 2013), however, no FDA approved targeted therapies are currently available for chondrosarcoma (Monga, Mani, Hirbe, & Milhem, 2020). Hence, the identification of novel targets for new treatment options remains urgent. Here, we found the expression of EZH2 was augmented in chondrosarcoma cell lines, which exerted a critical role in tumor progression. Over the past years, EZH2 inhibitors have been generated and tested. The compound, DZNep was shown to attenuate EZH2 protein expression and subsequently decreased H3K27me3, resulted in promoting cell death of chondrosarcoma cell line by apoptosis (Girard et al., 2014). In comparison to our results, selective EZH2 inhibitor, GSK343 and the FDA-approved tazemetostat were utilized to treat chondrosarcoma cell lines to examine their colony-forming ability. The inhibitory effects of colony formation of chondrosarcoma cell lines by GSK343 (5-10μM) (Supplementary Fig. 3) exhibited lower efficiency in contrast to the crizotinib (0.2-1μM) treatment *in vitro* (Fig. 6). It seems that the FDA-approved cMET inhibitors showed more portent retardation on chondrosarcoma tumorigenesis than inhibition of currently available EZH2 inhibitors.

Sulfatase can hydrolyze sulfate ester bonds of a broad range of substrates to remove 6-O-sulfate groups including HSPGs, which act as a co-receptor for various heparin-binding growth factors and cytokines subsequent to alter signaling pathways. Delineation of SULF1 expression in distinct epithelial cancer types has revealed complete loss or markedly decreased expression of SULF1 in various cancer cell lines (J. P. Lai, Chien, Strome, et al., 2004; J. P. Lai et al., 2008), suggesting that reduction of SULF1 is a general feature among epithelial cancers. Nevertheless, the role of SULF1 in chondrosarcoma is unclear. Therefore, enzymes that desulfate HSPGs could have a tumor suppressor effect via abolishment of ligand-receptor binding and downstream signaling. Mounting references showed the heparin-degrading endosulfatase SULF1 in cancers may contribute to affect cellular growth and survival signaling, tumor proliferation, migration, invasion and angiogenesis in vitro and in vivo (22). Prior studies revealed that the augmentation of 6-O-sulfation via three heparan sulfate (HS) 6-O-sulfotransferases (HS6ST1-3) may promote the cartilage tumor progression. Simultaneously, they also showed that expression of SULF1 didn’t exhibit significantly difference between benign tumor and distinct histological grade of chondrosarcoma (Waaijer et al., 2012). In contrast with our results, we observed the inhibition of SULF1 expression through histone 3 K27 trimethylation by EZH2 to attenuate the elimination of 6-O-sulfation in chondrosarcoma, resulting in the high level of 6-O-sulfation of HSPGs to enhance RTK signaling like cMET. Evidence has been demonstrated that downregulation of SULF1 in multiple cancers including ovarian, breast, gastric and hepatoma is orchestrated by DNA hypermethylation of CpG islands which located within promoter or exon 1A to prevent the transcription factors from binding to their DNA binding sites(J. P. Lai, Chien, Strome, et al., 2004; J. P. Lai, Chien, Moser, et al., 2004; Staub et al., 2007), concluding that epigenetic modulation was curial for SULF1 silencing in cancer cells.

Here, we showed that cMET phosphorylation was decreased in SULF1 overexpressed cells by using phospho-proteomic analysis of RTKs in vector and SULF1 ectopic expressed cell lines, suggesting that SULF1mediated the activity of RTKs. cMET has been reported to facilitate migration of chondrosarcoma cell lines upon HGF interaction (Tsou et al., 2013). Also, cMET signaling contributes to tumor survival, growth, angiogenesis and metastasis, and several cMET inhibitors were approved by FDA for cancer treatment. Prior investigation demonstrated that CD44, highly expressed in surface of chondrocytes and was a receptor for hyaluronan. Moreover, it binds to HGF and cMET to form complex to induce oncogenic signaling. Numerous post-translational modifications including N-and O-linked glycosylation, phosphorylation, sulfation and domain cleavage, which shapes the complication and function of CD44 protein family. For instance, the proinflammatory cytokine like tumor necrosis factor-α (TNF-α) was illustrated to change the inactive form of CD44 into active form via induction of the sulfation of CD44 (Maiti, Maki, & Johnson, 1998). Consistently, we showed that reduction of the sulfation of CD44 by SULF1 leading to inactivate the HGF, cMET, and CD44 tricomplex and repressed its downstream signaling.

Although there is no FDA approval targeted drug for chondrosarcoma, plenty of drugs are tested *in vivo* studies and clinical trials. Previously, various groups attempted to identify appropriate targets for prediction and therapies of chondrosarcoma by using diverse approaches. For instance, Nicolle *et al* indicated that integrated multi-omics classification to reveal the significance of the loss of expression of the 14q32 locus in defined malignancy on cartilage tumors. The isocitrate dehydrogenase (IDH) activating mutations driving a genome-wide hypermethylation in chondrosarcoma contributes to the activation of proliferative and glycolytic state, which was potentially driven by the hypoxia inducible factor (Nicolle et al., 2019). The other group uncovered that HIF-2α as a critical regulator for tumor growth, local invasion, and metastasis in chondrosarcoma by using weighted gene co-expression network analysis (WGCNA) (Kim et al., 2020). They also demonstrated that combination with small molecule HIF-2α inhibitor and chemotherapy agents synergistically promoted chondrosarcoma cell apoptosis and abrogates the malignant features of chondrosarcoma in mice.

Recently, Palubeckaite and colleagues have generated an alginate-based 3D cell culture model of chondrosarcoma to evaluate the treatment of chemotherapeutic drugs and target therapeutic agents including Sapanisertib (mTOR inhibitor) and AGI-519 (inhibitor of mutant IDH1). They reported that more resistant results of those treatment were observed in 3D compared to 2D cell culture system (Palubeckaite et al., 2020). Inhibition of mTOR alone in this 3D system would not have a higher effectiveness, and the improvement of therapeutic effects may occur by using combination treatment which could eliminate the remining cell population (Boehme, Schleicher, Traub, & Rolauffs, 2018). High expectations for effective treatment appeared while the characterization of specific driver mutations in the isocitrate dehydrogenase genes IDH1 and IDH2, and a specific inhibitor of mutant IDH1 was developed like AGI-5198. Yet, studies including using 3D culture model showed reduction of clonogenic capacity only after long-term treatment under high concentrations, while no benefit of treatment was detected at lower concentrations or shorter treatment times (L. Li et al., 2015; Suijker et al., 2015). Improved version of IDH mutation inhibitor to increase the metabolically stability, ivosidenib (AGI-120), was approved for chondrosarcoma clinical trial (Popovici-Muller et al., 2018) and revealed poor progression free survival with only one agent therapy in the retrospective investigation (van Maldegem et al., 2019). Suggesting that the limiting effects on the target therapies in chondrosarcoma thus far.

Tivantinib (ARQ-197) is a potent non-ATP competitive selective and oral c-Met small molecule inhibitor currently under clinical trial evaluation for liver and lung cancers (Michieli & Di Nicolantonio, 2013). Here, we also showed its potent inhibitory effect on phosphorylation of cMET and colony-forming ability in chondrosarcoma cell lines. In comparison to crizotinib, capmatinib was more specific for inhibition of MET (Vansteenkiste, Van De Kerkhove, Wauters, & Van Mol, 2019), yet chondrosarcoma seems to require higher dose of IC_50_. Crizotinib is a type Ia tyrosine kinase inhibitor that is approved for the treatment of ALK or ROS1-rearranged advanced non-small cell lung cancers (NSCLCs) (Gozdzik-Spychalska et al., 2014). In addition to its activity against ALK and ROS1, it has potent activity against cMET and low nanomolar potency in cell lines including our studies. It is not yet clear why crizotinib showed a better IC_50_ than capmatinib, one possibility might be due to the ability to target multiple RTKs. It is certainly worthy of pursuing further the underlying regulatory mechanisms.

We demonstrated that EZH2/ SULF1 axis mediates cMET in chondrosarcoma, and inhibition of cMET at low dose for long-term administration effectively alleviates tumor growth of chondrosarcoma and prolongs survival in mice. The effective therapeutic efficacy by crizotinib in chondrosarcaoma targeting EZH2.SULF1/ cMET shown in this report provides proof of concept evidence to use cMET as biomarker to select chondrosarcoma patients for treatment by the FDA-approved cMET inhibitors such as crizotinib.

## Material and method

### Cell culture

Human chondrosarcoma cell line, JJ012, was provided from Dr. Joel Block and maintained in Dulbecco’s modified Eagle’s medium (DMEM), F12, minimum essential medium (MEM), and fetal bovine serum (FBS) in a ratio of 40: 10: 40: 10%, supplemented with insulin, ascorbic acid and hydrocortisone. CH2879 was kindly provided from Drs. Antonio Llombart-Bosch were cultured in RPMI1640 contained 10% FBS. Patients-derived tissues were provided by Dr. Teng-Le Huang (CMUH IRB No.102-REC1-047) and isolated (Goldring, 1996). Normal primary chondrocytes were culture with DMEM/ F12 contain 1% P/S/F and 10% FBS.

### Real-time reverse transcription polymerase chain reaction (PCR)

Total RNA will be isolated using Trizol (Invitrogen) according to the manufacturer’s instructions. After total RNA isolation, cDNA was generated by SuperScript TM III reverse transcriptase (Invitrogen) and analyzed by real-time PCR (Roche, LightCycler® 480) with SYBR Green. All reaction products were normalized to the mRNA expression level of β-actin.

### Quantitative Chromatin immunoprecipitation (qChIP) assay and ChIP-seq

Chromatin immunoprecipitation (ChIP) assays were performed by EZ-ChIP kit (Upstate) according to the manufacturer’s instructions. Specific antibodies were used for immunoprecipitation in the ChIP assays. The immunoprecipitated DNA were subjected to real time PCR by Roche Cybr Green system according to the manufacturer’s instructions. For ChIP-seq analysis, the immunoprecipitated DNA was sequenced by Center for Genomic Medicine in NCKU (National Cheng Kung University) and sequenced data was further analyzed through NCKU bioinformatics center.

### Lentiviral infection

The lentiviral shRNA clones will be purchased from National RNAi Core Facility of Academic Sinica, Taiwan. Cells were infected with vector alone (pLK0.1) lentivirus or indicated shRNA lentivirus. Media containing lentivirus were collected and used to infect target cells supplemented with Polybrene (8 μg/ ml).

### Immunoblotting and immunoprecipitation

Cells were harvested on ice by using NETN (150 mM NaCl, 1 mM EDTA, 20 mM Tris-HCl, 0.5% NP40) buffer supplemented with protease inhibitors and lysates were incubated with antibodies for overnight at 4 °C. Immunocomplexes were precipitated with protein A/ G (Roche Applied Science) for 3h at 4 °C. The beads were washed four times with NETN buffer. Proteins were then eluted from the beads by boiling in sample buffer for 5 min and then analyzed by electrophoresis on SDS-polyacrylamide gels, and transferred to polyvinylidene difluoride (PVDF) membranes. Whole-cell lysates were analyzed with the following antibodies: pcMET (#3077 Cell Signaling), cMET (#8198 Cell Signaling), CD44v6 (BBA13 R&D), pERK 1/ 2 (GTX59568 #9101 Cell Signaling), ERK 1/ 2 #4695 Cell Signaling, H3K27me3 ab6002 Abcam, Histone 3 (GTX122148, #9715 Cell Signaling), ß-Actin (sc-47778 Santa Cruz Biotechnology), EZH2 (#3147, #5246 Cell Signaling) pP38 (#9211 Cell Signaling), P38 (sc-7149 Santa Cruz Biotechnology), SULF1 (ab32763 Abcam), 10E4 (#370255-S, amsbio). The image detection was analyzed by ChemiDoc™ Touch Imaging System (Bio-Rad) and Image Lab software (Bio-Rad).

### Proliferation assay

The proliferative ability of JJ012 or CH2879 cells with or without EZH2 shRNA were seeded at a density of 3×10^3^ cells or 5×10^3^ cells/ well in a 96-well tissue culture plate, respectively. Cells density were measured via Cell titer 96 one aqueous cell proliferation kits (MTS assay) after seeding for 48 h by ELISA reader at OD 490 nm. Each treatment was tested in quadruplicate. The same experiment was independently repeated three times. The data were evaluated against shRNA control using Student’s *t* test. The proliferation of vector and SULF1 ectopic expressed JJ012 and CH2879 stable clones were seeded 2×10^4^ cells and 1.1×10^5^ cells per well in 6-well plate, respectively. The cells were incubated for 24, 48 and 72 h. After emptied culture medium, 1 mg/ ml MTT (3-(4, 5-ciemthylthiazol-2-yl)-2, 5-diphenyltetrazolium bromide, Cat. No. M6494, Invitrogen) was added to each well for 1 ml and followed by incubated for 4 h in the dark under humidified conditions at 37°C and 5% CO_2_. Excess MTT was removed and then added 1 ml DMSO to each well. The plate was shacked for 5 min and they were detected with OD 570 nm by ELISA reader. For IC_50_, JJ012 or CH2879 cells were seeded at 7×10^4^ or 9×10^4^ cell density into 24 well, respectively. Cells were treated with different concentrations of cMET and EZH2 small molecular inhibitors for 48hr. The cell viability was determined by MTT assay. IC_50_ values were calculated by concentration-response curve fitting using a four-parameter analytical method via prism software.

### Migration assay

The migration assay was performed using the transwells (CORstar COR-3422 pore size, 8 μm) in a 24-well plate. The bottom chambers were filled with cultured media containing final concentrations of 10 % heat-inactivated FBS. JJ012 and CH2879 cells at a density of 1 × 10^4^ cells in serum free cultured media were added to upper chambers. After incubation at 37°C for 16 h, the membrane was fixed with methanol and stained with Geimsa then analyzed for the number of stained cells that had migrated to the opposite side of the membrane.

### Human phospho-RTK antibody array

Proteome Profiler Human phosphor-RTK array kit (ARY001B; R&D Systems, Minneapolis, MN) was used to detect potential activation of RTK signals by ectopic expressed SULF1 stable transfectants. All procedures were conducted according to the manufacturer’s instruction. Shortly, capture antibodies for specific RTKs were spotted in duplicate onto nitrocellulose membranes, provided by the kit. Suggested cell lysates were incubated with the array membrane at 4 °C overnight. To wash the unbound material, protein with phosphorylated tyrosine residues from the extracted cell lysates were bound to the capture antibodies of RTK and detected by a pan-phospho-tyrosine antibody conjugated with HRP. Finally, the binding signal was determined by chemiluminescent reagent and exposed to x-ray film.

### Establishment of stable cell lines

According to the type of vector, specific antibiotics were selected to generate the stable cells. G418 (1000 μg/ ml) were used to selected JJ012 stable cells; G418 (200 μg / ml) for CH2879 stable cells. All stable cells were selected from a single clone. G418 sulfate (Cat. No. Ant-gn-1, Invivogen). was purchased from InvivoGen Company.

### Flow cytometric analysis

For the analysis of sulfated glucosamine containing HSPG on cell membrane, 3×10^6^ of CH2879 and JJ012 stable cell lines were collected in 3% BSA and stained with 10E4 primary antibody (1:200 for 30 min; GTX20073), which recognizes native heparan sulfate containing the N-sulfated glucosamine moiety. After primary antibody staining, cells were further stained with Mouse IgM FITC (1:200) for 30 min at 4°C in the dark. The samples and data were analyzed by FACSVerse (BD) and its software BD FACSuite v1.0.6).

### Immunohistochemical staining and scoring of human bone sarcoma

Human bone sarcoma tissue array (B0481, B0481a) was purchased from US Biomax Inc and normal primary cartilage tissue array were collected from Dr. Teng-Le Huang (CMUH IRB No.102-REC1-047). Paraffin blocks were deparaffinized and hydrated through an ethanol series. The pressure cooker contains a retrieval solution (Tris-EDTA pH 9.0) and antigen retrieval was used by retrieval enzyme (ab970, abcam). Next, slides were treated with hydrogen peroxide and blocked by blocking buffer and then incubated with corresponding primary antibodies (pcMET 1:100, #3077 Cell Sinaling, cMET 1:100 #8198 Cell Sinaling, SULF1 1:100 ab32763, Abcam) overnight at 4 °C. The staining procedures were referred to manufactory’s instructions (Leica, RS-I-515-1). The Digital images of IHC-stained slides were obtained at 40x magnification (0.0625 μm^2^ per raw image pixel) using a whole slide scanner (Leica, Aperio CS2). Protein expression was ranked according to Histoscore (H-score) method. H-score was examined by a semi-quantitative evaluation of both the intensity of staining and percentage of positive cells (quantitative by Leica, image scope software, use Color Dconvolution v9 algorithm). The range of scores was from 0 to 250.

### Animal studies

All animal experiments were conducted according to animal welfare guidelines approved by China Medical University’s institutional Animal care and Use committee. For subcutaneous tumor model, 6-week-old female Balb/c nu/nu mice were inoculated 6×10^6^ vector and SULF1 CH2879 stable cells. Tumors were measured by vernier calipers every 4 days and the tumor volume was calculated using the following formula: (V)=π/ 6 × larger diameter × smaller diameter^2^. The mice were sacrificed after 45 days. For the orthotopic mice model, 6-week-old female Balb/c nu/nu mice were inoculated 5×10^5^ CH2879 cells ectopic expressed with luciferase. Cells were resuspended in 20µl PBS and injected into mouse tibial. After one-week tumor was formed and followed by treated with crizotinib via oral gavage 5 days per week. The tumor volume was measure by the IVIS system, and mice were weighed once a week. The mice survival rate was following the signs of either moribundity or mortality.

### Statistical analysis

Each experiment was repeated three times. Unless otherwise noted, data are presented as mean ± standard deviation. Statistical significance was determined by the Student t-test (two-tailed, unpaired). The statistical analysis for each plot is described in the figure legends. P < 0.05 was considered statistically significant.

### Antibody, primer sequence and commercial reagent

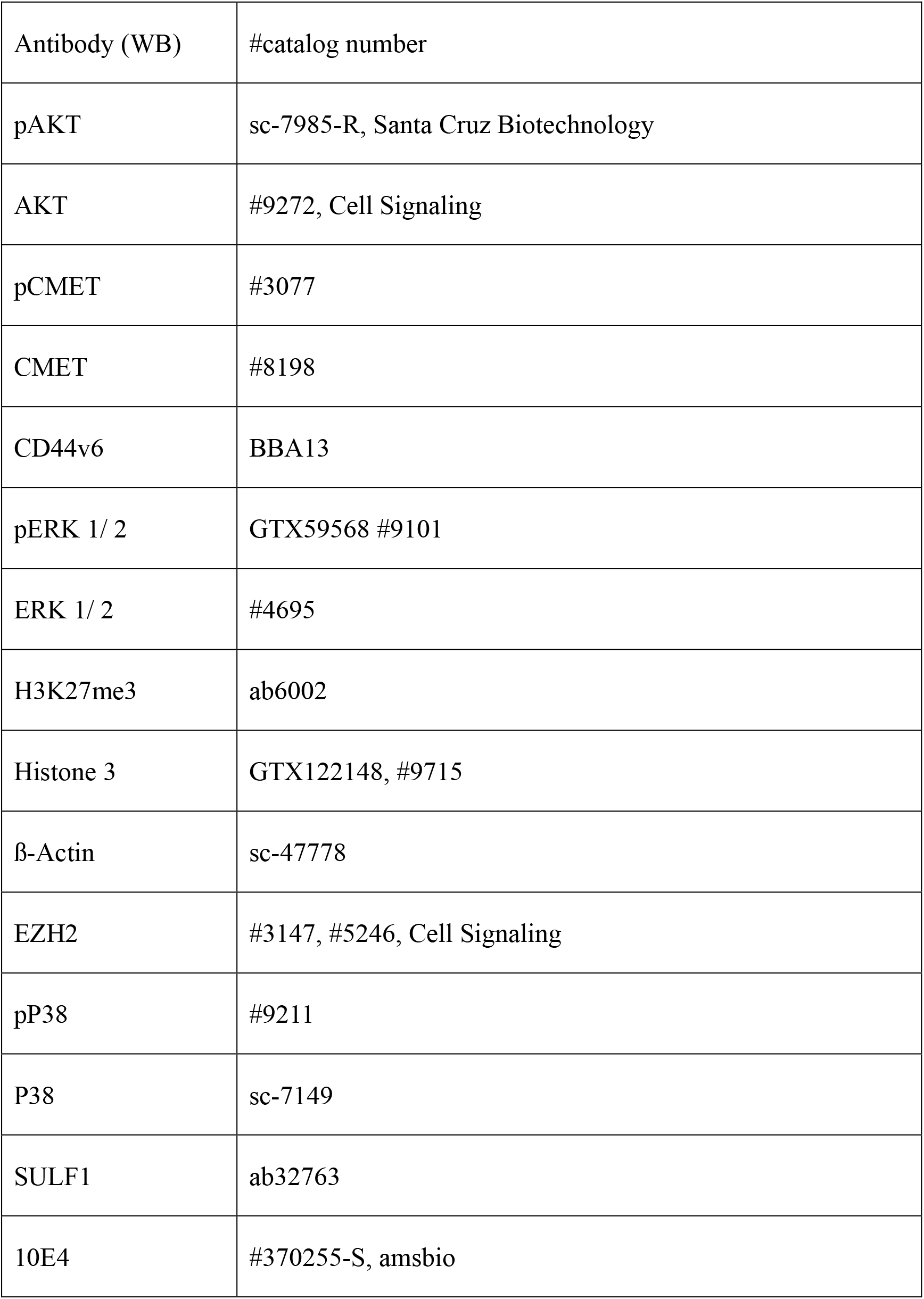
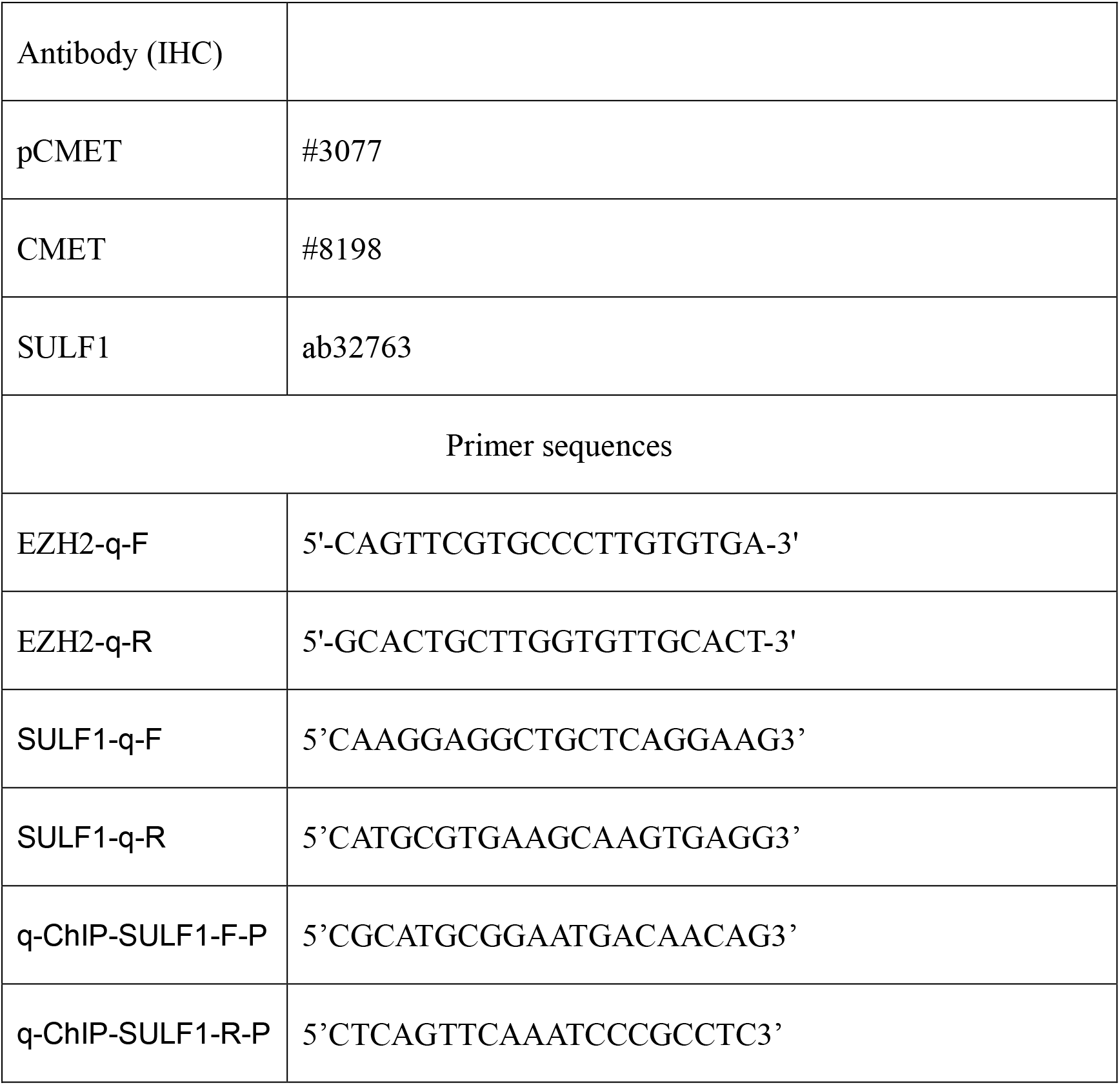

## Acknowledgments

We would like to acknowledge Dr. JA Block (Rush University Medical Center, Chicago, IL, USA), who provided us with the JJ012 cell line and Professor A Llombart Bosch (University of Valencia, Spain) for the CH2879 cell line. This study was supported by the following: Ministry of Science and Technology (MOST: 107-2320-B-039-003; To Y-H. C.; 108-2320-B-039-003; To Y-H. C.; 109-2320-B-039-012; To Y-H. C.); Drug Development Center, China Medical University from the Featured Areas Research Center program within the framework of the Higher Education Sprout Project by the Ministry of Education (MOE) in Taiwan (To Y-H. C.); China Medical University (CMU 107-TU-07; to Y-H. C; 107-S-36.). MOST 110-2639-B-039-001 -ASP (to M-C. H).

## Author contributions

Z.-S. L. designed and performed the experiments, analyzed data and wrote the manuscript; C.-C. C., Y.-C. L., C.-H. C., H.-C. L., Y.-Y. L., performed experiments and analyzed data; T.-L. H., and C.-H. T. provided patient tissue samples; T.-M. C., and C.-H. L. provided scientific input; M.-C. H., and Y.-H. C. supervised the entire project, designed the experiments, analyzed data and wrote the manuscript.

---

**SFig 1.**
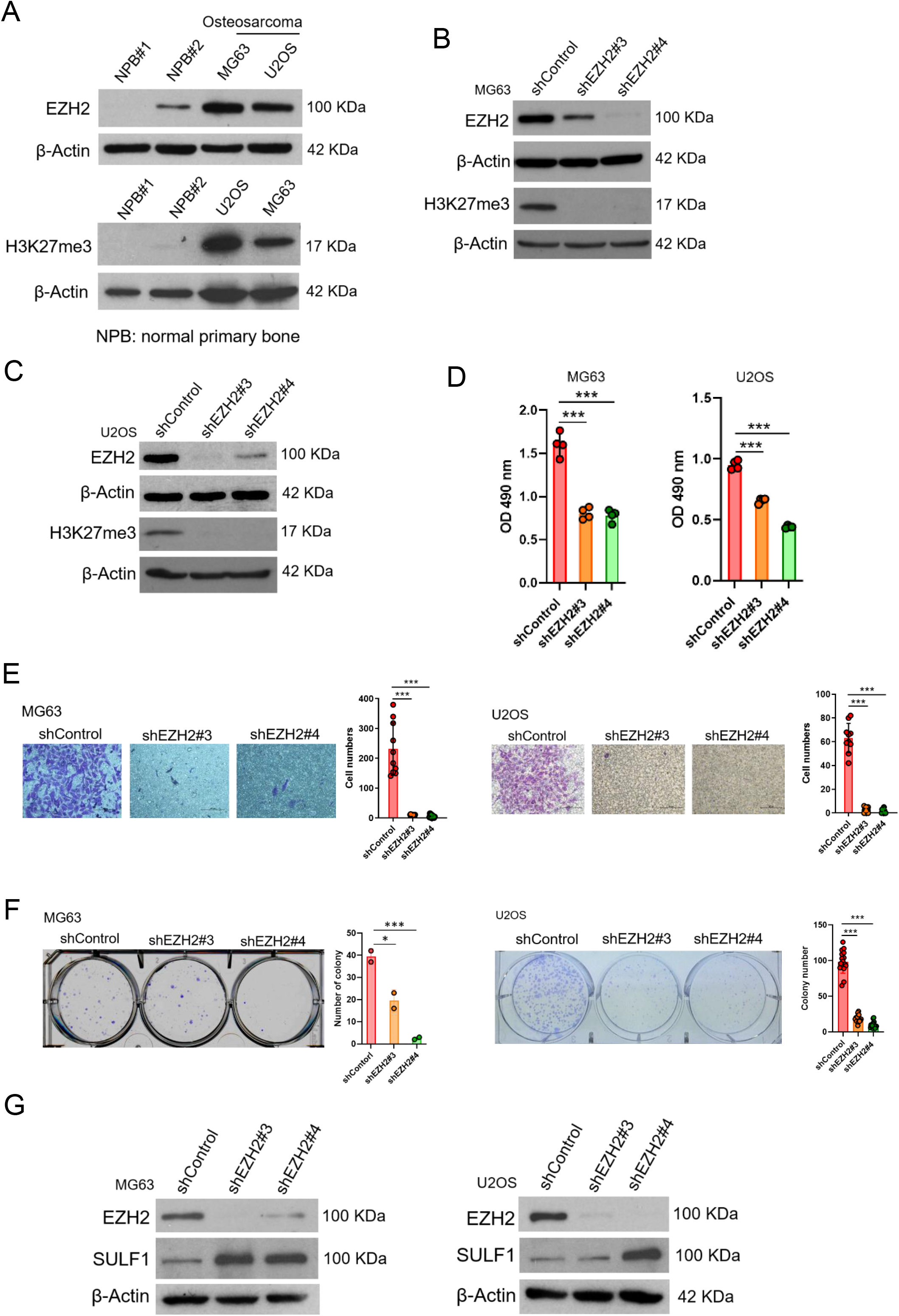

**SFig 2.**
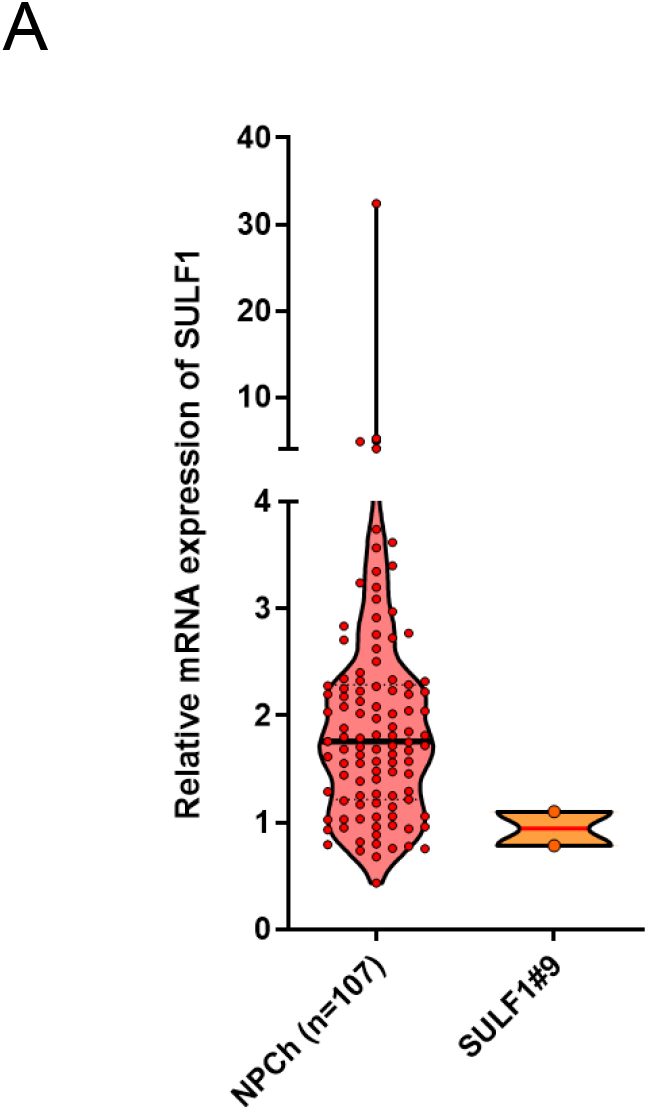

**SFig 3.**
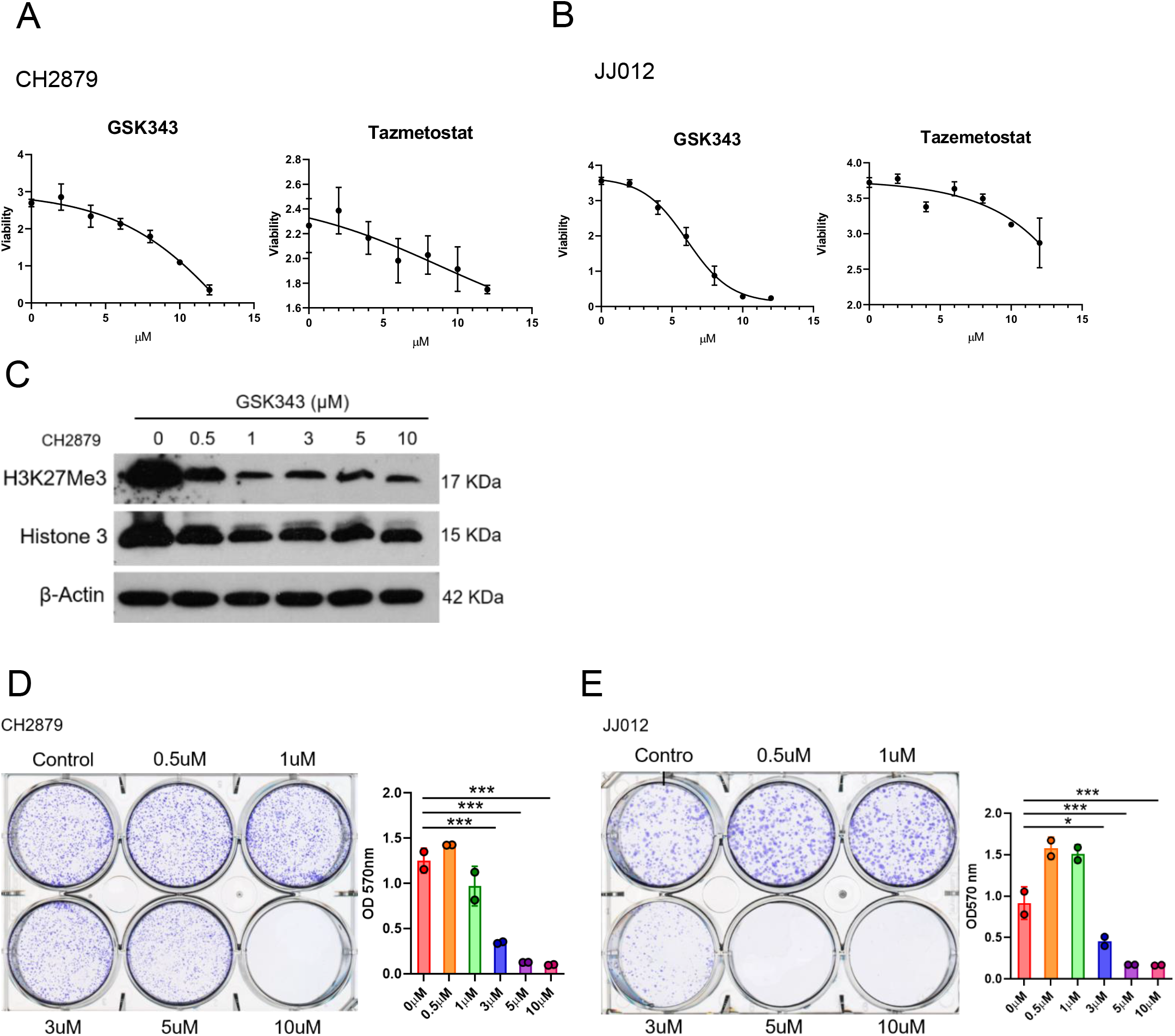

**SFig 4.**
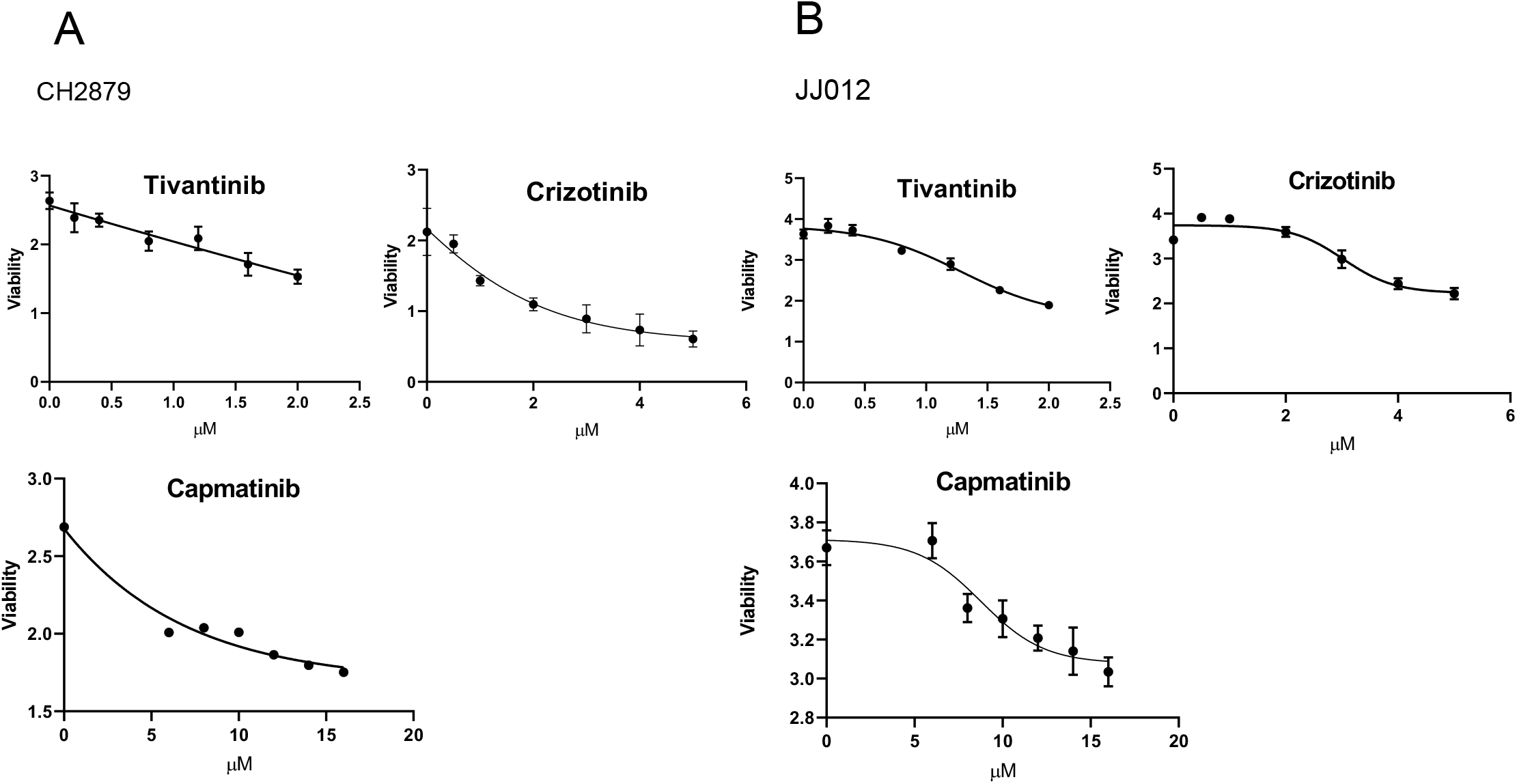

**SFig 5.**
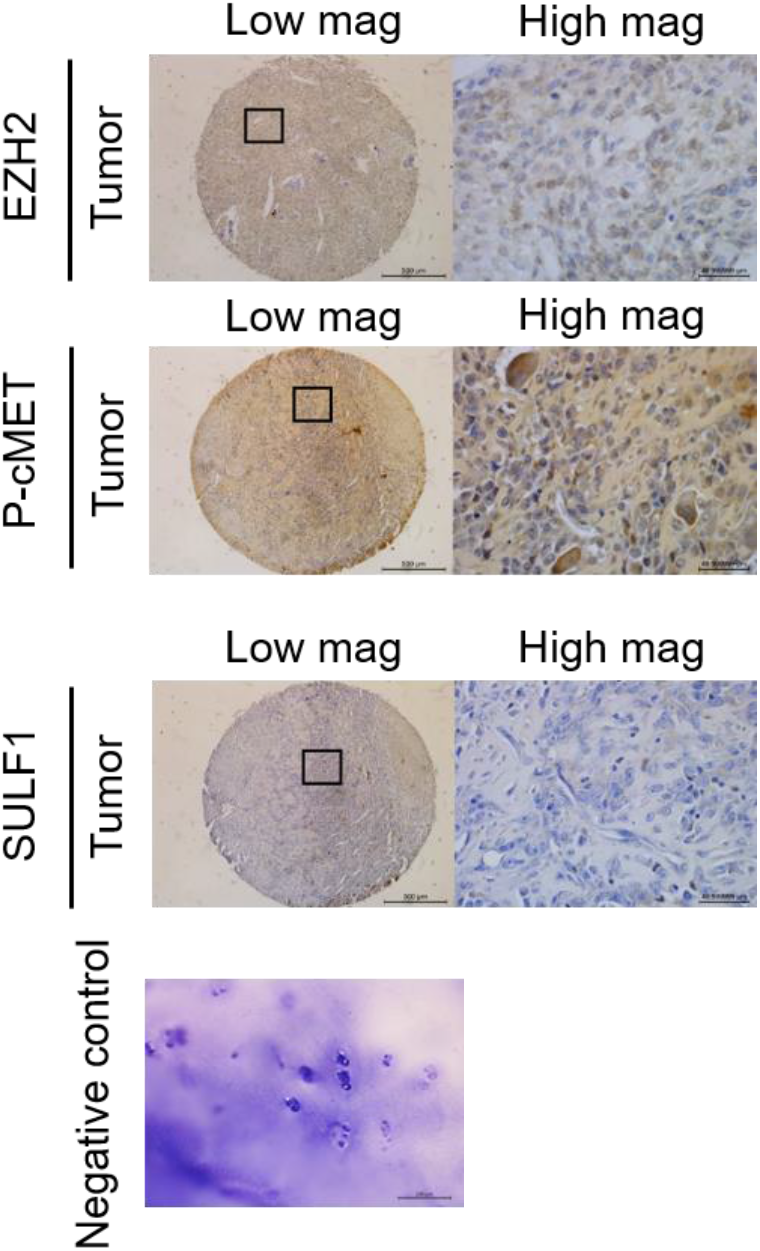

